# Aversion encoding and behavioral state modulation of lateral habenula neurons

**DOI:** 10.1101/2024.09.09.612004

**Authors:** Ioannis S. Zouridis, Lisa Schmors, Salvatore Lecca, Mauro Congiu, Manuel Mameli, Philipp Berens, Andrea Burgalossi

**Affiliations:** Institute of Neurobiology, Eberhard Karls University of Tübingen, Tübingen, Germany; Werner Reichardt Centre for Integrative Neuroscience, Tübingen, Germany; Graduate Training Centre of Neuroscience, International Max-Planck Research School (IMPRS), Tübingen, Germany; Hertie Institute for AI in Brain Health, University of Tübingen, Tübingen, Germany; Department of Fundamental Neuroscience, University of Lausanne, Lausanne, Switzerland; Inserm, UMR-S 839, 75005 Paris, France; Tübingen AI Center, University of Tübingen, Tübingen, Germany

**Keywords:** Lateral habenula, electrophysiology, juxtacellular, heterogeneity, diversity, cell types, clustering, Gaussian Mixtures Modeling, aversion, behavioral state

## Abstract

The lateral habenula (LHb) integrates aversive information to regulate motivated behaviors. Despite recent advances in identifying neuronal diversity at the molecular level, *in vivo* electrophysiological diversity of LHb neurons remains poorly understood. Understanding this diversity is essential for deciphering how information is processed in the LHb. To address this gap, we conducted *in vivo* electrophysiological recordings in mice and applied unsupervised clustering algorithm to analyze firing patterns. This analysis identified four distinct spontaneous firing patterns of LHb neurons, which were consistent across both anesthetized and awake states. To determine whether these firing patterns correlate with function, we recorded neuronal responses to foot shock stimulation in anesthetized mice and monitored spontaneous behavior in awake mice. We found that low-firing, bursting neurons were preferentially modulated by foot shocks in anesthetized mice and also tracked behavioral states in awake mice. Collectively, our findings indicate significant electrophysiological diversity among LHb neurons, which is associated with their modulation by aversive stimuli and behavioral state.

## 1 Introduction

The lateral habenula (LHb) is a bilateral epithalamic structure that integrates signals associated with aversive experiences to regulate motivated behaviors [1, 2]. The LHb was initially thought to be a homogenous brain structure, comprised almost exclusively of excitatory, glutamatergic neurons [3]. However, recent work has revealed a high degree of heterogeneity among LHb neurons with respect to their morphological, physiological, molecular, and connectivity properties [4, 5, 6, 7, 8]. Understanding this diversity is essential for deciphering how LHb integrates information to influence behavioral states and behavior [9].

Ex-vivo studies have classified LHb neurons into several morphological and electrophysiological subtypes [7, 8]. The LHb is organized into hodologically distinct medial and lateral territories, with the medial territory receiving projections from the bed nucleus of the stria terminalis and the lateral territory from the entopeduncular nucleus, each holding segregated projections to the ventral tegmental area and the rostromedial tegmental nucleus [10, 9, 11, 12]. The progress in omics technologies has confirmed this diversity at the molecular level [5, 6]. Moreover, transcriptionally defined LHb subtypes have been shown to have distinct connectivity, indicating a modular network architecture [6]. The existence of substantial diversity among LHb neurons is further supported by *in vivo* observations of diverse spiking patterns as well as variable functional properties; for example, different LHb units respond differently to aversive stimuli and engage differently in learning [13, 4]. However, the diversity of *in vivo* activity patterns and how they relate to the encoding of aversive stimuli and behavior remains poorly understood.

In this study, we aimed to fill this gap by clustering *in vivo* electrophysiological data to identify and quantify the diverse firing patterns of LHb neurons in both anesthetized and awake mice. Moreover, we related spontaneous firing patterns to responses to aversive stimuli and spontaneous behavioral state fluctuations. By resolving the diversity of LHb neurons and examining how this relates to their functional responses, we aim to provide a more accurate and detailed classification of LHb neurons and their modulation by aversive stimuli and behavioral states.

## 2 Results

### 2.1 LHb neurons spontaneously fire in four distinct patterns

To explore the diversity of spontaneous firing patterns of LHb neurons, we compiled datasets of *in vivo* juxtacellular recordings from anesthetized and awake preparations, and visualized the firing patterns in two-dimensional space using t-distributed Stochastic Neighbor Embedding (t-SNE, [14]; **Figure S2c,d**). To investigate the number and nature of spontaneous firing patterns of LHb neurons, we used Gaussian mixture model (GMM) clustering on each dataset separately. This analysis yielded four optimal clusters in each dataset, referred to as firing Type-1 through 4 thereafter (**Figure 1a,b**). To compare the firing pattern types of LHb neurons across awake and anesthetized states, we measured the cluster similarity by computing the cosine similarity (*S*_*C*_) and matching them using the Hungarian algorithm (see methods; **Figure 1c**, *S*_*C*_(1) = 0.76, *S*_*C*_(2) = 0.60, *S*_*C*_(3) = 0.82, *S*_*C*_(4) = 0.97). We identified corresponding firing pattern types with similar properties in both the anesthetized and awake datasets (**Figure 1d-h**). Although mean firing rates were considerably higher in the awake state (**Figure 1d, Figure S3a**), the direct correspondence of firing pattern types between the datasets indicates that these firing types are consistent across both states and not exclusive to one.

**Figure 1:**
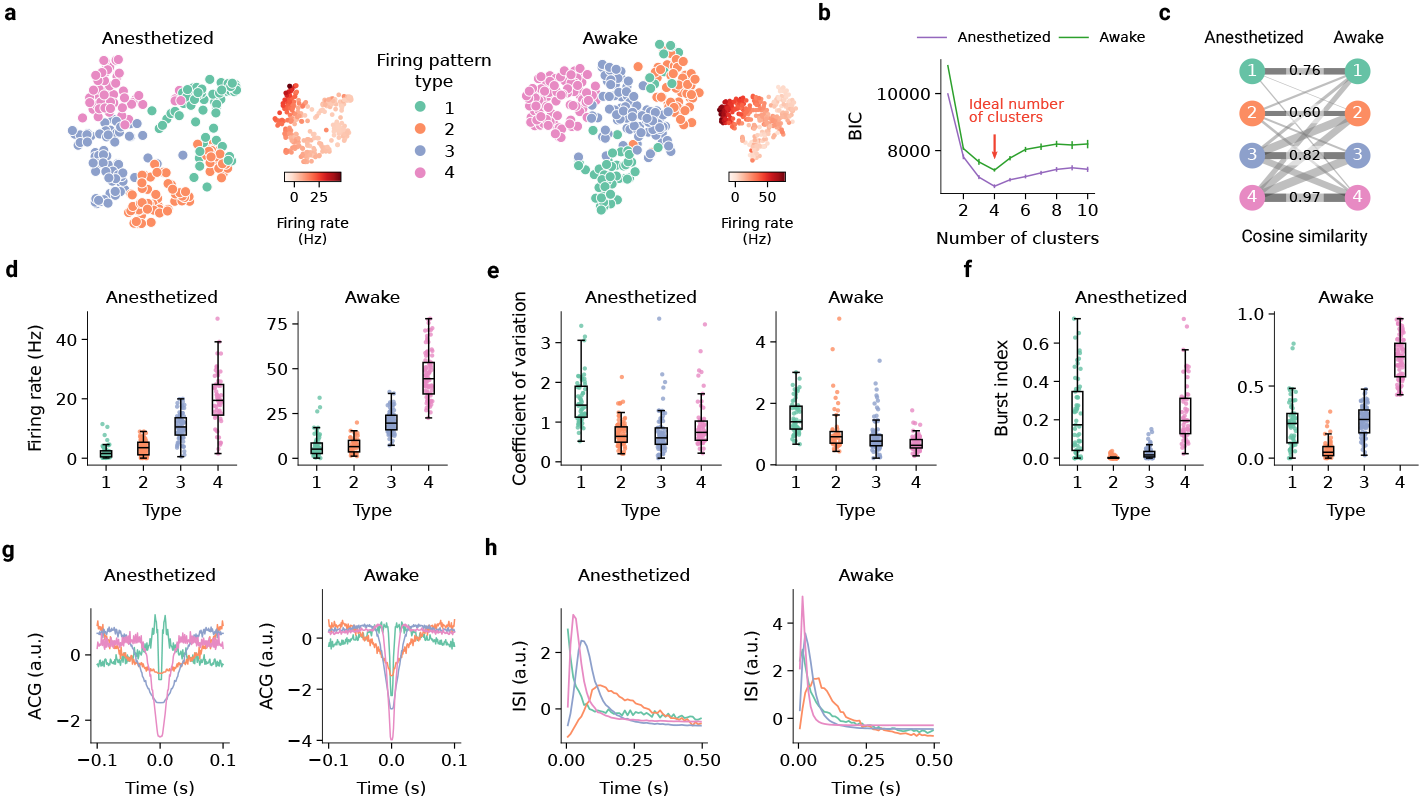
Gaussian Mixture Modeling of spontaneous firing features identifies four firing pattern types of LHb neurons. (**a**) t-SNE embedding based on 19 spontaneous firing features of LHb neurons recorded under anesthesia (left;n_neurons_ = 270) and awake recordings (right; n_neurons_ = 295). Each point represents a recording. Firing pattern types(color-coded) were determined using Gaussian Mixture Modeling (GMM) clustering using the same 19 features.Matching colors indicate the corresponding clusters across both conditions (see methods). The insets show t-SNE embedding colored by the mean spontaneous firing rate for each neuron. (**b**) Bayesian information criterion (BIC)computed for different numbers of clusters. Arrow indicates minimum BIC and the ideal number of clusters. For each number of clusters, the GMM was initialized randomly 100 times and the error bars show the confidence intervals across runs. (**c**) Clusters of anesthetized and awake recordings were matched using the Hungarian algorithm. Linewidths are proportional to the value of computed cosine similarity. (**d-f**) Mean firing rate (d), coefficient of variation(e), and burst index (f) for each firing pattern type under anesthesia (left) and awake (right). Number of recordings and conventions as in (a). (**g,h**) Mean autocorrelograms (g) and interspike interval distributions (h) for LHb neurons recorded under anesthesia (left) and awake recordings (right). Number of neurons and conventions as in (a).

Type-1 firing patterns were characterized by a low mean spontaneous firing rate (anesthetized: 2.07 ± 2.29; awake:7.15 ± 6.78; **Figure 1d**), high irregularity as indicated by a high coefficient of variation (anesthetized: 1.53 ± 0.58; awake: 1.57 ± 0.57; **Figure 1e**), and bursty activity as shown by a high mean burst index (anesthetized: 0.22 ± 0.20; awake: 0.24 ± 0.17; **Figure 1f**) and a peak at 5 ms in the autocorrelogram (ACG) and inter-spike interval (ISI) distributions (**Figure 1g,h**). Firing pattern Type-2 neurons exhibited a low mean spontaneous firing rate (anesthetized: 3.45 ± 2.47 Hz; awake: 7.37 ± 4.23 Hz; **Figure 1d**), a low burst index (anesthetized: 0.0038 ± 0.0078; awake: 0.06 ± 0.07; **Figure 1f**), and a tendency to fire regularly, as indicated by the ACG and ISI distributions (**Figure 1g,h**). Type-3 firing patterns showed a higher mean firing rate (anesthetized: 10.67 ± 4.61; awake: 20.18 ± 5.80; **Figure 1d**), a moderate burst index (anesthetized: 0.03 ± 0.03; awake: 0.26 ± 0.10; **Figure 1f**), and a low coefficient of variation (anesthetized: 0.74 ± 0.54; awake: 0.93 ± 0.54; **Figure 1e**). Type-4 firing patterns displayed a high firing rate (anesthetized: 19.57 ± 9.25; awake: 46.42 ± 13.32; **Figure 1d**), a high burst index (anesthetized: 0.24 ± 0.16; awake: 0.69 ± 0.15; **Figure 1f**), a low coefficient of variation (anesthetized: 0.95 ± 0.62; awake: 0.69 ± 0.24; **Figure 1e**), and an ISI distribution peaking at 25 ms (**Figure 1h**). We did not find an obvious anatomical organization of the different clusters within the LHb, based on our limited dataset of identified neurons **Figure S4**.

### 2.2 Type-1 firing pattern LHb neurons are preferentially modulated by aversive stimuli

LHb neurons are known to exhibit varied excitatory and inhibitory responses to aversive sensory stimuli, such as foot shocks (FS, [15, 13]; two example responses shown in **Figure 2a**). To investigate how different spontaneous firing pattern types relate to the response to aversive stimuli, we recorded neuronal responses to FS stimulation in anesthetized animals and computed a FS modulation index (FSMI; see methods). In a previous study, we demonstrated that the FSMI is negatively correlated with the spontaneous mean firing rate of LHb neurons [4]. Since neurons of different spontaneous firing types vary in their mean firing rates, we expected to observe differences also in their FS responses. Indeed, we observed varied responses across all four spontaneous firing types, with each type containing both FS-excited (FSMI > 0) and inhibited (FSMI *<* 0) neurons. For all firing pattern types, the mean FS responses were dominated by short-latency excitatory components (**Figure 2b,c**). Interestingly, neurons with spontaneous firing pattern Type-1 exhibited, on average, FS responses with a prominent long-latency excitatory component (response peaking at 210 ms; Figure **Figure 2b,c**). Further comparison of Type-1 against other spontaneous firing patterns revealed an on average higher FS modulation in Type-1 neurons (ANOVA, *p* < 0.005 n_neurons_ = 219; Type-1 vs. Type-2: mean difference = -0.205, 95% CI [-0.318, -0.091], *p* < 0.001; Type-1 vs. Type-3: mean difference = -0.313, 95% CI [-0.423 -0.203], *p* < 0.001; Type-1 vs. Type-4: mean difference = -0.262, 95% CI [-0.379 -0.144], *p* < 0.001; Figure **Figure 2d,e**). These results indicate that Type-1 neurons are comparatively more strongly and sustainedly modulated by FS stimulation. Further, FS modulation was positively correlated with burstiness (Pearson *r* = 0.29, *p* = 3.76 × 10^−5^) and negatively correlated with firing rate (Pearson *r* =− 0.36, *p* = 4.47 × 10^7^), both prominent in Type-1 neurons (**Figure S2e**).

**Figure 2:**
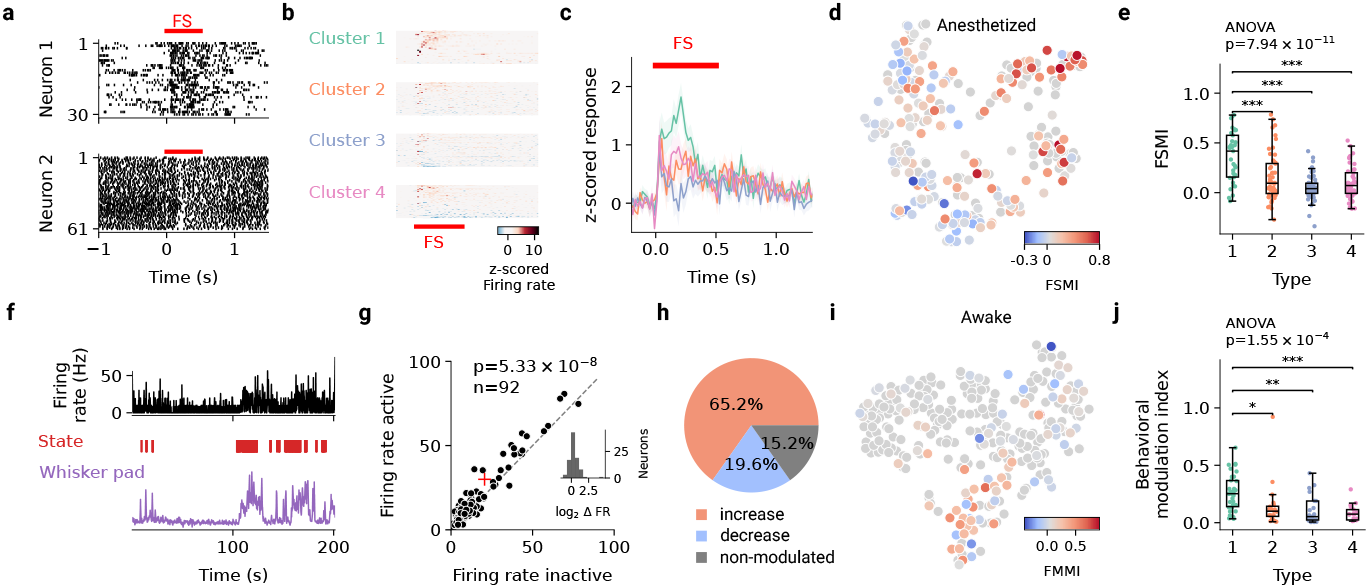
Sensory and behavioral state modulation of LHb neurons across different firing pattern types. (**a**) Responses to foot shock (FS) stimulation for two representative neurons in a spike raster plot. FS stimulation onset and duration indicated by a red line. (**b**) FS responses split by type. FS stimulation onset and duration indicated by a red line. (**c**) Mean FS responses of the four LHb neuronal types. FS stimulation onset and duration indicated by a red line.(**d**) t-SNE embedding of spontaneous firing patterns color-coded by the FS modulation index (n_neurons_ = 219; neurons without FS recordings are shown in gray). (**e**) Comparison of FS modulation index across different LHb neuronal types (ANOVA, n_neurons_ = 219, *p* = 7.94 × 10^−11^). Firing pattern Type-1 neurons were more strongly modulated in comparison to other types (Type-1 vs. Type-2: mean difference = -0.2049, 95% CI [-0.3183, -0.0914], *p* < 0.001; Type-1 vs. Type-3: mean difference = -0.3131, 95% CI [-0.4232, -0.2030], *p* < 0.001; Type-1 vs. Type-4: mean difference = -0.2615, 95% CI [-0.3788, -0.1441], *p* < 0.001; Type-2 vs. Type-3: mean difference = -0.1082, 95% CI [-0.2044, -0.0120], *p* = 0.0206). Significance bars with ∗ for *p* < 0.05, ∗ ∗ for *p* < 0.01, and ∗ ∗ ∗ for *p* < 0.001.(**f**) Representative recording of a LHb neuron along with orofacial videography. Top: firing rate; middle: behavioral state, red marks indicate epochs of ‘active’ state; bottom: whisker pad motion energy over time. (**g**) Effect of behavioral state on LHb neurons’ firing rate. *p* value denotes the result of a paired Wilcoxon signed-rank test, *n* = 92, mean: 20.58 Hz vs. 29.97 Hz, indicated with red cross. *Inset:* Histogram of firing rate fold-change in ‘active’ relative to ‘inactive’ behavioral states (Δ FR log_2_-ratio). (**h**) Summary of LHb neurons’ firing rate modulation by behavioral state, showing number and percentage of neurons with increases, decreases, or no change in activity. (**i**) t-SNE embedding of awake recordings color-coded by behavioral state modulation index (n_neurons_ = 92; neurons without facial motion recorded depicted in gray). (**j**) Behavioral state modulation index across different LHb firing patterns (ANOVA, n_neurons_ = 92, *p* = 1.55 × 10^−4^). Type-1 neurons were more strongly modulated in comparison to other types (Type-1 vs. Type-2: mean difference = -0.118, 95% CI [-0.229, -0.008], *p* = 0.031; Type-1 vs. Type-3: mean difference = -0.151, 95% CI [-0.259, -0.042], *p* = 0.003; Type-1 vs. Type-4: mean difference = -0.180, 95% CI [-0.296, -0.063], *p* <= 0.001). Significance bars as in (e).

### 2.3 Type-1 firing pattern LHb neurons are preferentially modulated by behavioral state

Recent work in awake head-fixed mice has demonstrated that spontaneous behavioral state fluctuations are accompanied by brain-wide modulations of neuronal activity [16, 17, 18, 19, 20]. Notably, while this activity modulation appears to be a brain-wide phenomenon, only a fraction of neurons are significantly modulated within each brain area, pointing to cell-type-specific mechanisms. To investigate whether and how the different LHb neuronal types are modulated by behavioral state, we acquired juxtacellular recordings along with orofacial videography in awake head-fixed mice. Using whisker-pad motion energy to index behavioral states [21, 17], we categorized the animal’s behavioral state into ‘active’ and ‘inactive’ (see methods; **Figure 2f**). On average, facial motion was associated with a significant net increase in neuronal activity (Wilcoxon test, n_neurons_ = 92, *p* = 5.33 × 10^−8^, 20.58 vs. 29.97 Hz; **Figure 2g**). However, individual LHb neurons were diversely modulated by spontaneous behavioral state fluctuations: the majority of neurons (60/92, 65.2%) showed a significant increase in mean firing rates during periods of ‘active’ behavioral state, while some exhibited reduced mean firing rates (18/92, 19.6%), or no significant modulation (14/92; 15.2%; **Figure 2h**; see Methods). Notably, these findings are consistent with LHb responses to FS, where most neurons were excited, while a minority were inhibited or non-responsive [13, 4].

To quantify the degree of behavioral state modulation, we computed a behavioral state modulation index (BSMI) for each neuron (see methods). We found that compared to other firing pattern types, Type-1 neurons were more strongly modulated by spontaneous behavior, as indicated by a comparatively higher BSMI (ANOVA, *p* < 0.005, n_neurons_ = 92; Type-1 vs. Type-2: mean difference = -0.118, 95% CI [-0.229,-0.008], *p* = 0.0306; Type-1 vs. Type-3: mean difference= -0.151, 95% CI [-0.259,-0.042], *p* = 0.0027; Type-1 vs. Type-4: mean difference = -0.180, 95% CI [-0.296,-0.063],*p* = 0.6 × 10^−3^; **Figure 2i,j** with results of Tukey’s HSD). These results indicate that LHb neurons are significantly modulated by spontaneous behavioral state fluctuations, with Type-1 neurons exhibiting, on average, the strongest modulation.

## 3 Discussion

In the present study, we investigated the physiological diversity of neurons in the mouse LHb, a brain region involved in processing aversive stimuli and regulating motivated behaviors. Previous work has revealed a high degree of electrophysiological and molecular diversity within LHb circuits; whether such diversity also applies to *in vivo* firing patterns has remained largely unexplored. By applying an unsupervised clustering approach to *in vivo* electrophysiological data, we identified four distinct spontaneous firing pattern types in LHb neurons. Notably, these firing pattern types were consistently observed in both anesthetized and awake states, indicating that this diversity might be imposed by structural determinants. Indeed, our observations are consistent with previous ex-vivo studies, which identified four distinct firing pattern types in the rodent LHb [8], and transcriptomic studies [6], which also identified four molecularly-defined LHb cell types, each with its own connectivity [6].

We note, however, that despite the above correspondences, our observations are based on unsupervised clustering of firing patterns. Hence, whether the four firing pattern types are discrete or rather map onto a continuum of neuronal features remains to be investigated. Moreover, whether the firing pattern type (i.e., cluster affiliation) of neurons is fixed (i.e., pre-determined by, e.g., intrinsic cellular properties and/or synaptic inputs) or dynamically modulated (e.g., by behavioral state and/or learning) remains to be investigated. We speculate that the diversity in firing patterns observed *in vivo* might correlate to the known diversity of intrinsic neuronal properties, morphologies, and input connectivity described ex-vivo [7, 8]. Indeed, the tendency of neurons to fire spike bursts and input-output connectivity [10, 9, 11, 12] are anatomically organized within the LHb. Future work will be required for testing this hypothesis and for resolving possible structure-function relationships within LHb circuits.

Spontaneous fluctuations in behavioral state modulate neuronal activity throughout the brain, yet only a small fraction of neurons in each brain structure are strongly modulated [16, 17, 18, 19, 20]. Indeed, in line with previous work, we found that only a minority of neurons within the LHb were particularly responsive to aversive stimuli in anesthetized mice and modulated by behavioral state in awake mice. Interestingly, under both anesthetized and awake conditions, these modulated neurons were preferentially recruited from Type-1 firing patterns. Compared to the other clusters, these neurons also displayed a higher tendency to fire spike bursts. Since spike bursts have been linked to plasticity and excitability [22, 23], our data indicate that this minority Type-1 LHb neurons might play a fundamental role in information encoding within the LHb. This hypothesis is indeed consistent with a large body of previous work, which demonstrated that minorities of ‘excitable & plastic’ cells dominate information encoding and processing in different brain areas like neocortex [24], hippocampus [25], and amygdala [26]. Neuronal spike bursts are also particularly efficient at overcoming synaptic failures and transmitting information to downstream targets [23]. Indeed, bursts in LHb neurons efficiently inhibit reward-related downstream targets and have been shown to play an important role in diseases including depression. [27] . Hence, Type-1 neurons might also play a dominant role in pathophysiological LHb states. We speculate that modulating the activity of Type-1 neurons may be a particularly powerful approach for modulating LHb computations as demonstrated, e.g., in the hippocampus, neocortex and amygdala, where artificial modulation of ‘active minorities’ has been shown to be sufficient for modifying memory content and for driving behavior [28, 29, 30]. Whether Type-1 LHb neurons can be preferentially tagged with immediate early gene expression as, e.g., in cortico-hippocampal and amygdala networks [31, 32] remains to be investigated.

In summary, our data provide insights into the diversity of LHb *in vivo* firing patterns and point to low-firing, bursty (Type-1) neurons playing a unique role in information encoding within LHb circuits.

## Acknowledgments

This work was supported by the Werner Reichardt Centre for Integrative Neuroscience (EXC 307), the Eberhard-Karls University of Tübingen, the DFG grants BU 3126/2-1 and BU 3126/3-1 (A.B.), the CRC 1233 ‘Robust Vision’ (TP 13, project number: 276693517; P.B.), the Swiss National Science Foundation 31003A-175549 (M.M.), the Swiss State Secretariat for Education, Research and Innovation SVEN (M.M.) and the Hertie Foundation. We thank Fabio Monteiro and Kathrin Maite Fisher for their excellent assistance in histology experiments.

## Author contributions

**Conceptualization**: I.S.Z., L.S., P.B., A.B.; **Data Curation**: I.S.Z. annotate & produce metadata, data preprocessing, maintain research data, maintain software code; L.S. data preprocessing, create & maintain software code; M.M. contributed metadata production; **Formal Analysis**: I.S.Z. data processing, statistical analysis; L.S. data processing, statistical analysis, computational models; **Funding Acquisition**: P.B. and A.B.; **Investigation**: I.S.Z. electrophysiological and behavioral recordings; M.C. and S.L. contributed electrophysiology data; **Methodology**: I.S.Z. electrophysiology, behavior, anatomy experiments; L.S. creation of computational models; **Project Administration**: I.S.Z., A.B., and L.S.; **Resources**: P.B. and A.B.; **Software**: L.S. software development for statistical analysis and computational models; I.S.Z. software for data preprocessing; **Supervision**: P.B., A.B., M.M.; **Validation**: I.S.Z. and L.S.; **Visualization**:L.S. designed and produced figures, I.S.Z. contributed designing figures; **Writing – Original Draft**: I.S.Z. and L.S.; **Writing – Review & Editing**: All authors.

## Data and code availability

Data and code will be made available upon submission.

## 4 Materials and methods

### 4.1 Experimental animals

Experiments were performed on adult male C57BL/6J mice (>8 weeks old, 20–30 g body weight; Charles River, Sulzfeld, Germany). The mice were housed under a 12-hour light cycle and were provided with unrestricted access to food and water. All experimental procedures were approved by local authorities and were performed according to guidelines of the respective local ethics committee: the canton of Vaud Cantonal Veterinary Office Committee for Animal Experimentation (Switzerland), in compliance with the Swiss National Institutional Guidelines on animal experimentation; the Regierungspräsidium Tübingen (Germany) in compliance with the German Animal Welfare Act (TierSchG) and the Animal Welfare Laboratory Animal Ordinance (TierSchVersV).

### 4.2 *In vivo* electrophysiological recordings

#### 4.2.1 *In vivo* electrophysiological recordings in anesthetized mice

*In vivo* electrophysiological recordings of LHb neurons in mice have been compiled from previous studies [13, 15, 4]. The procedures and datasets are described in detail in [4]. No additional recordings in anesthetized preparations have been conducted for this study. Briefly, recordings include juxtacellular recordings of spontaneous activity of LHb neurons in mice anesthetized with either a ketamine-xylazine mixture or isoflurane (recording duration: 220.90±89.05 s, ± mean SD). In a subset of mice (n=64), foot shock (FS) stimulation (1 mA amplitude, 0.5 s stimulus duration, repeated every 5 s, with on average 69 ± 44 trials per neuron) was applied to the contralateral hind limb. Comparative analysis indicated that neuronal firing properties were comparable between ketamine-xylazine and isoflurane-anesthetized mice (**Figure S1**), allowing us to combine recordings obtained under different anesthetics into a single dataset.

#### 4.2.2 Histological determination of the neurons’ anatomical location

For a subset of recorded and labeled neurons (part of the dataset acquired under ketamine-xylazine anesthesia in [4]), the anatomical location of the juxtacellularly labeled neuron was determined. The brain tissue was processed essentially as previously described in [33]. Briefly, serial tissue sections (70 *µ*m thickness; coronal sectioning plane) were obtained using a vibrating blade microtome (VT1200, Leica, Germany), stained with a fluorescent streptavidin conjugate (1:1000, Cat#S11225, Thermo Fisher Scientific) and DAPI (1:1000, Cat#D1306, Thermo Fisher Scientific), and examined using epifluorescence microscopy (Axio Imager, Carl Zeiss, Germany).

#### 4.2.3 *In vivo* juxtacellular recordings and orofacial videography in awake head-fixed mice

##### Surgical procedures

Surgery, habituation, and recordings in awake head-fixed mice were performed as previously described [34, 17]. Briefly, mice (n=10) were anesthetized with a ketamine-xylazine mixture (80-100 and 10 mg/kg, respectively, i.p.), fixed on a stereotaxic frame (Narishige, Tokyo, Japan), and a custom-made head-post was attached to the skull using a light-curing adhesive (Optibond Universal, Kerr) and dental acrylic (Paladur, Heraeus Kulzer, Hanau, Germany). For targeting the LHb, a craniotomy was performed at the stereotactic coordinates: 1.5 mm posterior and 0.5 mm lateral to bregma. During the intervals between recording sessions, the craniotomy site was covered with a silicon sealant (Kwik-Cast, WPI).

##### *In vivo* juxtacellular recording and labeling

A neurobiotin-filled [1.5–2% Neurobiotin, SP-1120, Vector Laboratories in ‘intracellular-like’ solution (in mM): 135 K-gluconate, 10 HEPES, 10 Na2-phosphocreatine, 4 KCl, 4 MgATP, and 0.3 Na3GTP, osmolarity: 280–310 mOsm] glass pipette electrode [(Cat#1403547, Hilgenberg, Malsfeld, Germany; pulled to a 7-9 MOhm resistance with a micropipette puller (Cat#P-1000, Sutter Instrument, Novato, CA, USA)] was lowered into the LHb (coordinates: 2.3-2.9 mm ventral to the brain surface). Juxtacellular voltage signals were acquired using an ELC-03XS amplifier (NPI Electronic, Tamm, Germany) and digitized at 25 kHz (Power1401-3, Spike2 v8.02; CED, Cambridge, UK). Upon establishing juxtacellular configuration with a neuron, spontaneous spiking activity was monitored (mean recording duration: 228.11 s). When feasible, once the recording was completed, neurons were labeled by delivering repetitive, square current pulses (200 ms pulse duration, 50% duty charge) [35].

##### Histological verification of the recording location

To confirm the recording location in the LHb, animals were overdosed with pentobarbital (300 mg/kg, i.p.), transcardially perfused (0.1 M phosphate buffer followed by a 4% paraformaldehyde 0.1 phosphate buffer solution), and dissected. The brain tissue was processed essentially as described in methods 4.2.2. Recordings from a session were included in the analysis only if labeled cells or electrode tracks were confirmed to be within the LHb.

##### Orofacial videography

The mouse’s left whisker pad was illuminated with infrared light (M850L3, Thorlabs), video was recorded using an externally triggered camera (DMK33UX265, TheImagingSource), and trigger pulses were co-registered with electrophysiology data for post-hoc temporal alignment (see methods 4.2.3).

### 4.3 Quantification and statistical analysis

Data analysis was performed using custom-written code based on packages such as numpy (v. 1.22.3), sklearn (v. 1.0.2), matplotlib (v. 3.5.2), seaborn (v. 0.13.2), and pandas (v. 1.4.2). All statistical analyses were conducted using the Python libraries scipy (v. 1.8.0) and statsmodels (v. 0.13.2). To compare more than two groups, we used ANOVA with *α* = 0.05 and Tukey’s HSD post-hoc test. In total, we included 270 recordings in anesthetized animals from Congiu et al. (2023) [4], 88 of which were originally recorded in [13] and 33 in [15]. Our dataset in awake, head-fixed mice consisted of 295 recordings.

#### Electrophysiology data analysis

The analysis of juxtacellular electrophysiological data was performed as previously described (e.g., [33]). Specifically, spikes were isolated from continuous voltage traces using a threshold-based approach and visual inspection. For each neuron, we computed the mean firing rate, the interspike interval (ISI) distribution (range 0–0.5 s), the coefficient of variation (CV; calculated as the ratio of the standard deviation of the ISI over the mean ISI [36]), the burst index (calculated as the percentage of ISIs shorter than 25 ms), and the autocorrelogram (ACG; range ± 0.1 s) based on spontaneous activity. Temporal features of the ISI distributions and ACGs were extracted with sparse principal component analysis (sPCA; [37]) using the first 8 principal components (PCs; **Figure S2a,b**).

#### Clustering

We fit each data set with a Mixture of Gaussians model [38] using the following 19 features: mean FR, CV, burst index, first 8 PCs of the ISI distribution, and first 8 PCs of the ACG (**Figure S2a–d**). We applied the Bayesian Information Criterion (BIC; [39]) to determine the optimal number of clusters: *BIC* =− 2 log[*L*] + *M* log[*N*], with *L* as the likelihood function for the model, and *M* as the number of parameters in the model. The BIC naturally implements Occam’s razor by favoring simpler models (fewer clusters) unless there is strong evidence in the data for more complex models (more clusters). To avoid local minima, we initialized the GMM algorithm 100 times per candidate cluster number (error bars in **Figure 1b**) and selected the number of clusters with the smallest mean BIC, which resulted in four clusters for both data sets. To match clusters from the anesthetized preparation (A) with clusters from the awake preparation (B), we used the cosine similarity *S*_*C*_ [40] to measure the similarity between cluster centroids: *S*_*C*_(cluster_*A*_, cluster_*B*_) 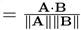 We applied the Hungarian algorithm [41, 42] to find the optimal matching between clusters from the two datasets. It optimally matches clusters by minimizing the total dissimilarity min 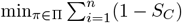, where *π* is a permutation of cluster indices and Π is the set of all possible combinations. We further filtered matches based on a similarity threshold of 0.6, ensuring that only cluster pairs with sufficient similarity were included.

#### Analysis of foot shock responses

To assess the responses of neurons to foot shock stimuli, we constructed peristimulus time histograms ( −0.2 to +1.3 s with respect to stimulus onset; bin width, 20 ms; **Figure 2c**). To quantify the strength of foot shock modulation, we computed a foot shock modulation index (FSMI) [34], defined as:

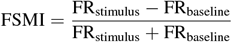

where FR_baseline_ represents the mean firing rate during 1 s preceding the stimulus onset, FR_stimulus_ represents the mean firing rate during stimulus delivery. Neurons with firing rate *<* 1 Hz were excluded because calculating the index for these neurons is not feasible due to them potentially generating fewer than one spike per stimulation duration.

#### Videography data analysis

To assess the behavioral state, we used gross facial motion as an index of activity. Whisker pad motion energy was extracted using Facemap [16] as detailed in [17]. To partition behavior into states, we defined ‘active’ epochs as those with whisker pad motion energy above 80th percentile and ‘inactive’ epochs below 20th percentile. For every neuron, we tested the firing rate modulation in ‘active’ versus ‘inactive’ behavioral states using an unpaired Wilcoxon signed-rank test at a 0.05 significance level. Neurons were split into neurons that show a significant firing rate increase or decrease or no significant modulation in response to behavioral state. To quantify the strength of modulation, we computed a behavioral state modulation index (BSMI), defined as:

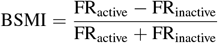

where FR_active_ represents the mean firing rate during epochs of ‘active’ behavioral state, FR_inactive_ represents the mean firing rate during epochs of ‘inactive’ behavioral state.

## Supplementary Information

**Figure S1:**
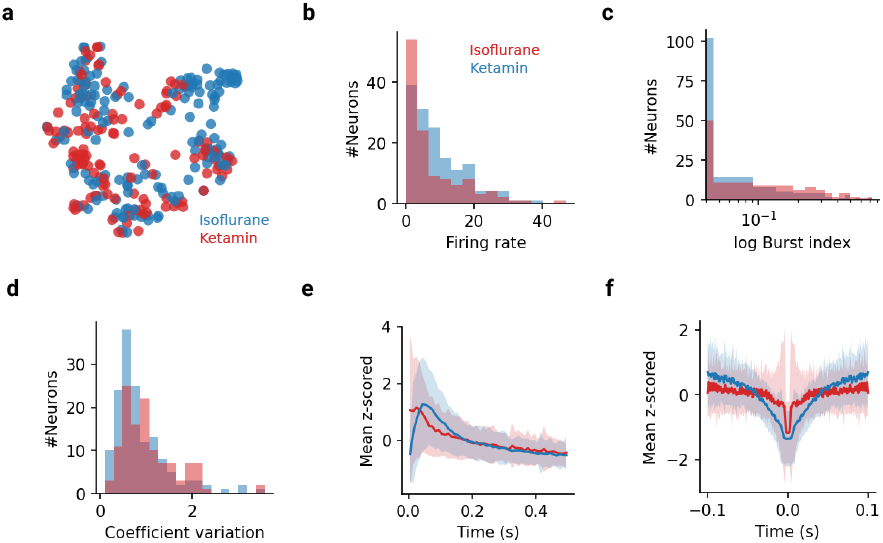
Electrophysiological properties of LHb neurons recorded under ketamine-xylazine or isoflurane anesthesia. (**a**) t-SNE embedding of electrophysiological properties of LHb neurons recorded under ketamine (red; n_neurons_ = 149) or isoflurane (red; n_neurons_ = 121) anesthesia. Each point represents a recording. (**b**) Mean firing rate distribution of neurons recorded under ketamine or isoflurane anesthesia. (**c**) Burst index distribution of neurons recorded under ketamine or isoflurane anesthesia. While NMDA antagonism has been shown to abolish bursting in LHb neurons, we observed Type-1 patterns under ketamine-xylazine anesthesia. As expected, bursting was more prominent under isoflurane anesthesia. (**d**) Coefficient of variation of neurons recorded under ketamine-xylazine or isoflurane anesthesia. (**e**) Mean interspike interval distribution of neurons computed for neurons from ketamine-xylazine and isoflurane conditions. Shaded area indicates ± 1 SD. (**f**) Mean autocorrelograms computed for neurons from ketamine-xylazine and isoflurane anesthesia. Shaded area indicates ± 1 SD.

**Figure S2:**
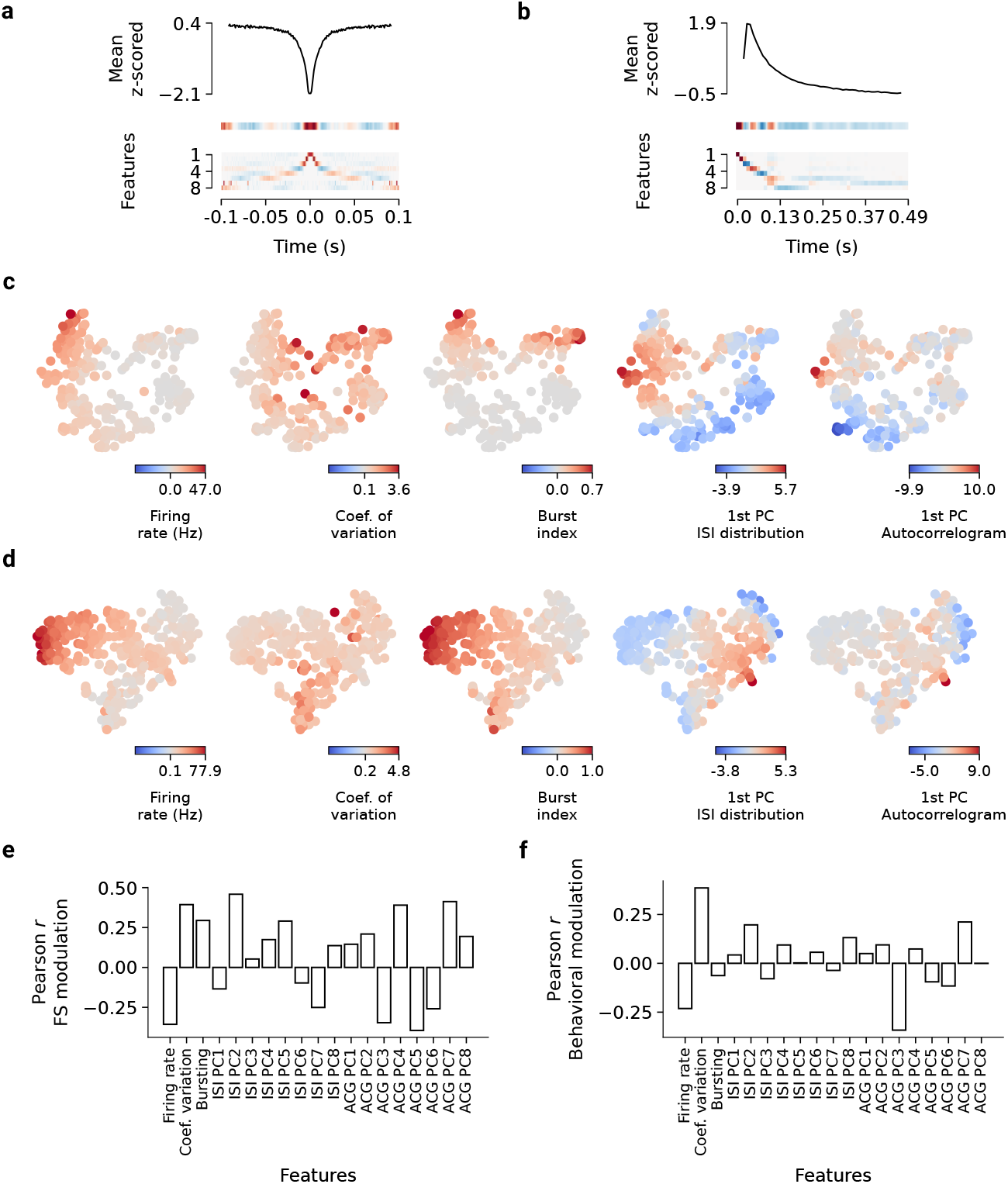
Distributions of features and their correlations across LHb neurons in anesthetized and awake conditions. (**a**) ACG feature extraction using sparse Principal Components Analysis (sPCA). *Top:* mean z-scored ACG for all neurons (black; n_neurons_ = 565). *Bottom:* all 8 sPCA components. (**b**) Same as in (a) but for interspike-interval. (**c**) t-SNE embedding of electrophysiological properties of LHb neurons recorded under anesthesia. Neurons are color-coded by the respective feature, from left to right: firing rate, coefficient of variation, burst index, principle component for ISI, and principal component for ACG (only the first principle component is shown here). (**d**) Same as in (c) but for neurons recorded under anesthesia. (**e**) Pearson correlation coefficient (*r*) between foot shock modulation and features used for clustering. Refer to panels (a) and (b) for the interpretation of sPCA components. (**f**) Same as in (e) but for modulation by behavioral state.

**Figure S3:**
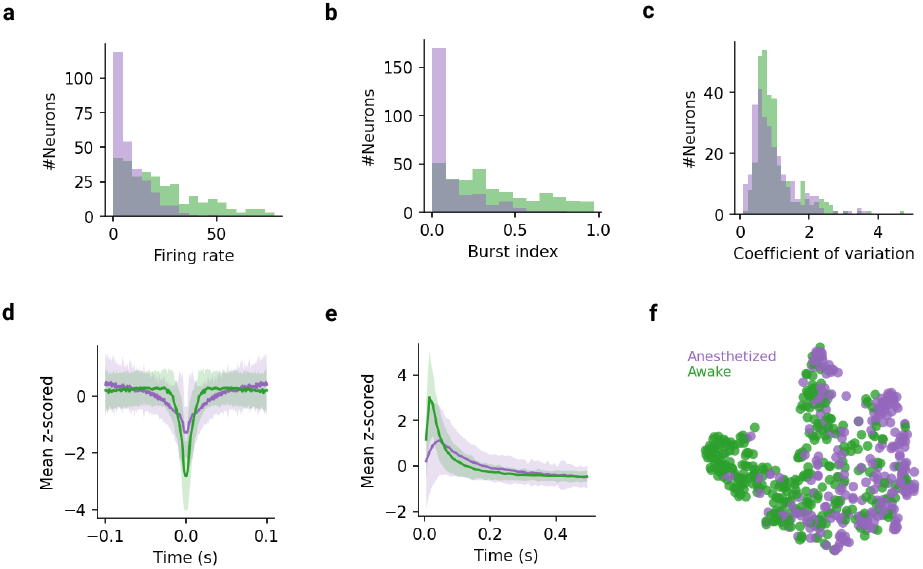
Electrophysiological properties of LHb neurons recorded under anesthesia and awake conditions. (**a-c**) Distributions of firing rate (a), burst index (b), and coefficient of variation (c) of LHb neurons recorded under anesthesia (purple; n_neurons_ = 270) and awake (green; n_neurons_ = 295) conditions. (**d-e**) Autocorrelograms (d) and interspike interval distributions (e) computed for LHb neurons recorded under anesthesia (purple) and awake (green) conditions. Lines indicate the mean, shaded areas indicate ±1 SD. (**f**) t-SNE embedding of electrophysiological properties of LHb neurons recorded under anesthesia (purple; n_neurons_ = 270) and awake conditions (green; n_neurons_ = 295). Each point represents a recording.

**Figure S4:**
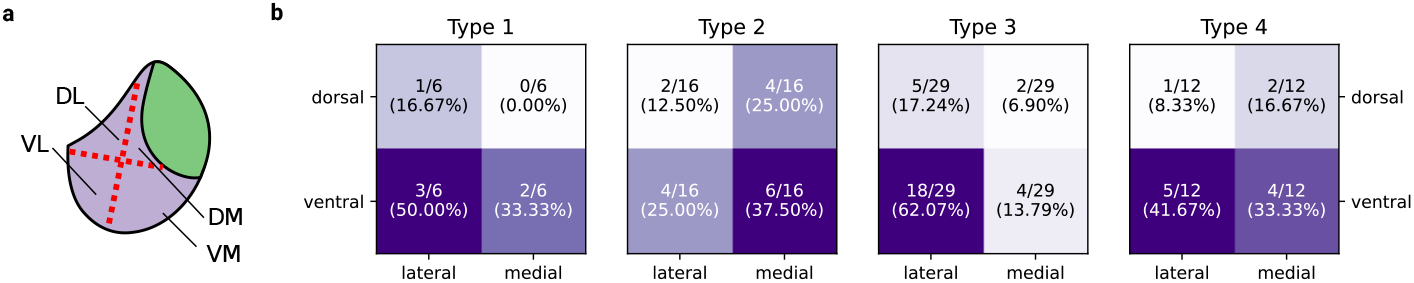
Topographical distribution of firing pattern types in the LHb. (**a**) Schematic representation of the partitioning of LHb in four quartiles, namely dorsomedial (DM), dorsolateral (DL), ventromedial (VM) and ventrolateral (VL). Anatomical orientation: Dorsal, up; lateral, left. LHb, magenta; Medial habenula, green. (**b**) Topographical distribution of firing patterns in quartiles. Firing pattern Type-1, dorsolateral (DL): 1/6, dorsomedial (DM): 0/6, ventrolateral (VL): 3/6, ventromedial (VM): 2/6; Firing pattern Type-2: DL: 2/16, DM: 4/16, VL: 4/16, VM: 6/16; Firing pattern Type-3: DL: 5/29, DM: 2/29, VL: 18/29, VM: 4/29; Type-4: DL: 1/12, DM: 2/12, VL: 5/12, VM: 4/12

